# The amino acid substitution affects cellular response to mistranslation

**DOI:** 10.1101/2021.05.12.443880

**Authors:** Matthew D. Berg, Yanrui Zhu, Bianca Y. Ruiz, Raphaël Loll-Krippleber, Joshua Isaacson, Bryan-Joseph San Luis, Julie Genereaux, Charles Boone, Judit Villén, Grant W. Brown, Christopher J. Brandl

## Abstract

Mistranslation, the mis-incorporation of an amino acid not specified by the “standard” genetic code, occurs in all organisms. tRNA variants that increase mistranslation arise spontaneously and engineered tRNAs can achieve mistranslation frequencies approaching 10% in yeast and bacteria. Interestingly, human genomes contain tRNA variants with the potential to mistranslate. Cells cope with increased mistranslation through multiple mechanisms, though high levels cause proteotoxic stress. The goal of this study was to compare the genetic interactions and the impact on transcriptome and cellular growth of two tRNA variants that mistranslate at a similar frequency but create different amino acid substitutions in *Saccharomyces cerevisiae.* One tRNA variant inserts alanine at proline codons whereas the other inserts serine for arginine. Both tRNAs decreased growth rate, with the effect being greater for arginine to serine than for proline to alanine. The tRNA that substituted serine for arginine resulted in a heat shock response. In contrast, heat shock response was minimal for proline to alanine substitution. Further demonstrating the significance of the amino acid substitution, transcriptome analysis identified unique up- and downregulated genes in response to each mistranslating tRNA. Number and extent of negative synthetic genetic interactions also differed depending upon type of mistranslation. Based on the unique responses observed for these mistranslating tRNAs, we predict that the potential of mistranslation to exacerbate diseases caused by proteotoxic stress depends on the tRNA variant. Furthermore, based on their unique transcriptomes and genetic interactions, different naturally occurring mistranslating tRNAs have the potential to negatively influence specific diseases.

## INTRODUCTION

Mistranslation, the incorporation of an amino acid not specified by the “standard” genetic code, occurs with frequencies varying from 10^−2^to 10^−5^, depending on the codon (reviewed in Joshi *et al.* 2019). Mistranslation increases in specific environmental conditions [reviewed in Mohler and Ibba (2017)] and due to mutations in the translation machinery, including in tRNA encoding genes [reviewed in Berg and Brandl (2020)]. In particular, tRNA aminoacylation plays a key role in maintaining translation fidelity, since the ribosome validates the codon-anticodon pairing and not the amino acid (Chapeville *et al.* 1962; Ogle *et al.* 2002; Loveland *et al.* 2017). Aminoacyl-tRNA synthetases (aaRS) recognize their cognate tRNAs using structural elements and nucleotides within the tRNA called identity elements (Rich and RajBhandary 1976; de Duve 1988; Giegé *et al.* 1998). For many tRNAs, recognition is determined by the anticodon. This is not the case for tRNA^Ser^ and tRNA^Ala^ (Giegé *et al.* 1998). For tRNA^Ser^, the main identity element is the long variable arm positioned 3’ of the anticodon stem (Asahara *et al.* 1994; Biou *et al.* 1994; Himeno *et al.* 1997). Therefore, anticodon mutations in tRNA^Ser^ encoding genes lead to mis-incorporation of serine at non-serine codons (Geslain *et al.* 2010; Berg *et al.* 2017, 2019b; Zimmerman *et al.* 2018). The main identity element for tRNA^Ala^ is a G3:U70 base pair in the acceptor stem (Imura *et al.* 1969; Hou and Schimmel 1988, 1989). Addition of this element to non-alanine tRNAs results in mis-aminoacylation and mistranslation of alanine at non-alanine codons (McClain and Foss 1988; Francklyn and Schimmel 1989; Hoffman *et al.* 2017a; Lant *et al.* 2017).

Mutations in tRNAs that cause mistranslation arise spontaneously. Early studies identified intergenic suppressors that change the meaning of the genetic code (Stadler and Yanofsky 1959; Yanofsky and Crawford 1959; Crawford and Yanofsky 1959; Benzer and Champe 1962; Gorini and Beckwith 1966). These include tRNA variants that lead to glycine mistranslation at arginine or cysteine codons in *Escherichia coli* (Carbon *et al.* 1966; Jones *et al.* 1966; Gupta and Khorana 1966) and a tRNA^Tyr^ variant in yeast that suppresses stop codons (Goodman *et al.* 1977). Similar tRNA variants that likely mistranslate are found in human populations. In a sample of 84 individuals, we identified six variants that create G3:U70 base pairs in non-alanine tRNAs and 14 anticodon variants that alter tRNA decoding identity (Berg *et al.* 2019a).

Multiple copies of the tRNA genes and overlapping isoacceptors allow cells to tolerate the mistranslation caused by tRNA variants; for example, *Saccharomyces cerevisiae* contain approximately 275 tRNA encoding loci. In addition, cells have protein quality control mechanisms to cope with mis-made proteins resulting from mistranslation. These include the ubiquitin-proteasome system, autophagy, induction of the heat shock and unfolded protein responses, and the organization of aggregates into inclusion bodies (reviewed in Hoffman *et al.* 2017b). When mistranslation reaches a threshold, protein quality control mechanisms no longer protect the cell. Ruan *et al.* (2008) demonstrated that *E. coli* tolerate mistranslation frequencies of approximately 10%. Our work suggests that yeast cells remain viable with mistranslation approaching 8% (Berg *et al.* 2019b). Based on the prevalence of naturally occurring tRNA variants that potentially mistranslate, we hypothesize that mistranslation in humans could modulate disease, especially diseases characterized by loss of proteostasis (see for example Lee *et al.* 2006; Liu *et al.* 2014; Lu *et al.* 2014 and reviewed in Lant *et al.* 2019).

The abundance of naturally occurring tRNA variation, the relationship between tRNA variation and mistranslation and the association of mistranslation with disease led us to investigate the genetic interactions and the impact on the transcriptome and cellular growth of different mistranslating tRNA variants in yeast. Previously, we found that repression of yeast growth caused by tRNA^Ser^ variants that mis-incorporate serine at proline codons correlated with the frequency of mistranslation (Berg *et al.* 2019b). However, when investigating tRNAs that mistranslate serine at a range of non-serine codons, Zimmerman *et al.* (2018) found that decreased growth did not simply correlate with mistranslation frequency. In this study, we compare the impact of two tRNAs variants that mistranslate at similar frequency but substitute different amino acids. Using variants that mistranslate either alanine at proline codons or serine at arginine codons, we find differences in growth rate, heat shock response, transcriptomes, and genetic interactions for each of the two tRNA variants. We suggest that the nature of the amino acid substitution is important in determining the impact of human tRNA variants on diseases caused by protein mis-folding, and that different tRNA variants have the potential to negatively influence distinct diseases, aside from those characterized by protein mis-folding.

## MATERIALS AND METHODS

### Yeast strains and growth

BY4741 (*MATa his3∆0 leu2∆0 met15∆0 ura3∆0*; Brachmann *et al.* 1998), BY4742 (*MATα his3∆0 leu2∆0 lys2∆0 ura3∆0*; Brachmann *et al.* 1998) and Y7092 (SGA starting strain, *MATα can1∆::STE2pr-SpHIS5 lyp1∆ his3∆1 leu2∆0 ura3∆0 met15∆0*) strains are derivatives of S288c. Y7092 was a kind gift from Dr. Brenda Andrews (University of Toronto). Strains from the temperature sensitive collection are derived from the wild-type *MATa* haploid yeast strain BY4741 and described in Costanzo *et al.* (2016). Strains containing mistranslating tRNAs were made by integrating genes encoding tRNA^Pro^_G3:U70_ (CY8612) or tRNA^Ser^_UCU, G26A_ (CY8614) into Y7092 at the *HO* locus using the constructs described below and selecting for the *natNT2* marker. The control strain (CY8611) was made by integrating only the *natNT2* marker at the *HO* locus. Transformants were selected on YPD media containing 100 µg/mL nourseothricin-dihydrogen sulfate (clonNAT) and integration was verified by PCR.

Yeast strains were grown in yeast peptone media containing 2% glucose (YPD) or synthetic media supplemented with nitrogenous bases and amino acids at 30°. Growth curves were generated by diluting saturated cultures to OD_600_ equal to 0.1 and incubating at 30°. Growth curves were performed either in YPD, YPD containing 4% ethanol, YP containing 2% galactose, synthetic complete media containing ammonium sulfate as the nitrogen source and either 2% glucose or 2% galactose as the carbon source, synthetic complete media containing monosodium glutamate as the nitrogen source and 2% glucose as the carbon source or minimal media containing ammonium sulfate as the nitrogen source, 2% glucose, adenine, histidine, leucine, tryptophan, lysine and methionine. OD_600_ was measured every 15 min using a BioTek Epoch 2 microplate spectrophotometer for 24 hr. Doubling time was calculated using the R package “growthcurver” (Sprouffske and Wagner 2016).

### DNA constructs

Constructs to integrate mistranslating tRNAs at the *HO* locus were created using synthetic DNA containing 200 bp up and downstream of the *HO* translational start as previously described in Zhu *et al.* (2020; Figure S1; Life Technologies). The construct was cloned into pGEM®-T Easy (Promega Corp.) as a *Not*I fragment to create pCB4386. The *natNT2* marker from pFA6-natNT2 was PCR amplified using primers UK9789/UK9790 (Table S1) and cloned into pCB4386 as an *Eco*RI fragment to generate the control SGA integrating vector (pCB4394). The gene encoding tRNA^Pro^_G3:U70_ was moved as a *Hin*dIII fragment from pCB2948 (Hoffman *et al.* 2017a) into pCB4394 to create pCB4396. The gene encoding tRNA^Ser^_UCU_, G26A was PCR amplified from pCB4224 (Berg *et al.* 2019b) using primers UG5953/VB2609 and cloned as *Hin*dIII fragments into pCB4394 to create pCB4398.

*URA3*-containing centromeric plasmids expressing tRNA^Ser^ (pCB3076), tRNA^Pro^_UGG G3:U70_ (pCB2948) and tRNA^Ser^_UCU, G26A_ (pCB4301) are described in Berg *et al.* (2017), Hoffman *et al.* (2017) and Berg *et al.* (2019). The centromeric plasmid containing HSE-eGFP was kindly provided by Onn Brandman (Stanford University; Brandman *et al.* 2012).

### Mass spectrometry

Liquid chromatography tandem mass spectrometry to identify mistranslation was performed on five biological replicates of the control strain (CY8611) and strains containing one of the mistranslating tRNAs (CY8612 or CY8614). Starter cultures of each strain were grown to saturation in YPD, diluted to an OD_600_ of 0.1 in the same media and grown to an OD_600_ of ~ 0.8 at 30°. Preparation of cell lysates, protein reduction and alkylation were performed as described in Berg *et al.* (2019b). Robotic purification and digestion of proteins into peptides were performed on the KingFisher™ Flex using LysC and the R2-P1 method as described in Leutert *et al.* (2019).

Peptides were analyzed on a hybrid quadrupole orbitrap mass spectrometer (Orbitrap Exploris 480; Thermo Fisher Scientific) equipped with an Easy1200 nanoLC system (Thermo Fisher Scientific). Peptide samples were resuspended in 4% acetonitrile, 3% formic acid and loaded onto a 100 μm ID × 3 cm precolumn packed with Reprosil C18 3 μm beads (Dr. Maisch GmbH), and separated by reverse‐phase chromatography on a 100 μm ID × 30 cm analytical column packed with Reprosil C18 1.9 μm beads (Dr. Maisch GmbH) housed into a column heater set at 50°.

Peptides were separated using a gradient of 5-30% acetonitrile in 0.125% formic acid at 400 nL/min over 95 min, with a total 120 minute acquisition time. The mass spectrometer was operated in data-dependent acquisition mode with a defined cycle time of 3 seconds. For each cycle one full mass spectrometry scan was acquired from 350 to 1200 m/z at 120,000 resolution with a fill target of 3E6 ions and automated calculation of injection time. The most abundant ions from the full MS scan were selected for fragmentation using 2 m/z precursor isolation window and beam‐type collisional‐activation dissociation (HCD) with 30% normalized collision energy. MS/MS spectra were acquired at 15,000 resolution by setting the AGC target to standard and injection time to automated mode. Fragmented precursors were dynamically excluded from selection for 60 seconds.

MS/MS spectra were searched against the *S. cerevisiae* protein sequence database (downloaded from the Saccharomyces Genome Database resource in 2014) using Comet (release 2015.01; Eng *et al.* 2013). The precursor mass tolerance was set to 50 ppm. Constant modification of cysteine carbamidomethylation (57.0215 Da) and variable modification of methionine oxidation (15.9949 Da) were used for all searches. Variable modification of proline to alanine (−26.0157 Da) or arginine to serine (−69.0691 Da) were used for the respective mistranslating tRNAs. A maximum of two of each variable modification were allowed per peptide. Search results were filtered to a 1% false discovery rate at the peptide spectrum match level using Percolator (Käll *et al.* 2007). The mistranslation frequency was calculated using the unique mistranslated peptides for which the non-mistranslated sibling peptide was also observed. The frequency is defined as the counts of mistranslated peptides, where alanine was inserted for proline or serine inserted for arginine, divided by the counts of all peptides containing proline or arginine, respectively, and expressed as a percentage.

### Synthetic genetic array analysis and validation

The SGA assay was performed as described by Tong *et al.* (2001) with minor modifications. Strains CY8611 (*HO::natMX*), CY8612 (*HO::tRNA^Pro^_G3:U70_-natMX*) and CY8614 (*HO∷tRNA^Ser^_UCU, G26A_-natMX*) were mated with a yeast temperature sensitive collection (Ben-Aroya *et al.* 2008; Li *et al.* 2011; Kofoed *et al.* 2015; Costanzo *et al.* 2016) in quadruplicate 1536 colony array format using a BioMatrix (S&P Robotics Inc.) automated pinning system. In this format, each allele of the temperature sensitive collection is present in biological quadruplicate on the plate. Double mutants were selected on YPD plates containing 200 mg/L G418 and 100 mg/L clonNAT. Diploids were sporulated on enriched sporulation media and *MATa* haploid double-mutants selected using standard SGA media. To identify genetic interactions, double mutants were pinned onto double mutant selection SGA medium and grown at 30° for 5 days. Images were taken every 24 hours to determine colony size computationally. SGATools (Wagih *et al.* 2013) was used to determine genetic interaction scores using a multiplicative model. Double mutant strains with average interaction score less than −0.2 and Benjamini-Hochberg corrected *p-*value less than 0.05 were considered as potential negative genetic interactions.

Double mutants that were identified as negative genetic interactions from the screen were validated by re-creating the double mutant strain, starting from the single mutant haploid strains, using the SGA approach. Double mutant strains were grown in liquid media to saturation, cell densities were normalized, and cultures were spotted on SGA media. The temperature sensitive mutant crossed with the control strain CY8611 and the mistranslating tRNA strain crossed with a control *his3∆* strain were also spotted to determine fitness of the single mutants. Intensity of each spot was measured with ImageJ (Schneider *et al.* 2012). Expected double mutant growth was calculated based on the growth of the single mutants and compared to the experimental growth of the double mutant. Double mutants that grew more slowly than expected were considered validated negative genetic interactions. Raw and validated data can be found in Supplemental File S2.

### RNA Preparation, Sequencing and Analysis

Wild-type yeast strain BY4742 containing either an empty *URA3* plasmid (control), pCB2948 (tRNA^Pro^_G3:U70_) or pCB4301 (tRNA^Ser^_UCU, G26A_) were grown to stationary phase in media lacking uracil. Strains were diluted in the same media to an OD_600_ of 5.0 × 10^−3^ and grown to a final OD_600_ of 2.0. Cells were harvested and RNA extracted with MasterPure™ Yeast RNA Purification Kit (Epicentre). Samples were vacuum dried and sent to Genewiz (South Plainfield, NJ) for total RNA sequencing.

Stranded Illumina TruSeq cDNA libraries with poly dT enrichment were prepared from high quality total RNA (RIN > 8). Libraries were sequenced on an Illumina HiSeq yielding between 30.2 – 39.5 million 150 bp paired end reads per sample.

FASTQ files were analyzed using a customized bioinformatics workflow. Adapter sequences and low quality bases were trimmed using the default settings of Trimmomatic (Bolger *et al.* 2014). Sequence quality was analyzed using FastQC (http://www.bioinformatics.babraham.ac.uk/projects/fastqc/). Reads were aligned to the *S. cerevisiae* S288C reference genome (Engel *et al.* 2014; release R64-2-1_20150113) using STAR (Dobin *et al.* 2013). Only uniquely mapping reads were counted. Read counts for each gene were summarized using featureCounts (Liao *et al.* 2014). Differential expression analysis was performed using the DESeq2 R package (Love *et al.* 2014) with a Benjamini-Hochberg adjusted *p*-value cut off of ≤ 0.05. Analysis script can be found in Supplemental File S3. The data have been deposited in NCBI’s Gene Expression Omnibus (Edgar *et al.* 2002) and are accessible through GEO Series accession number GSE174145.

### GO Term Analysis, Genetic Interaction Network Creation and SAFE Analysis

Gene Ontology (GO) term analysis was performed using the GO term finder tool (http://go.princeton.edu/) using a P-value cut-off of 0.01 after applying Bonferroni correction. Terms were filtered with REVIGO (Supek *et al.* 2011). Networks were constructed using Cytoscape 3.7 (Shannon *et al.* 2003).

### Heat shock assay

Yeast strains containing the *HSE-GFP* reporter (Brandman *et al.* 2012) and a mistranslating tRNA variant were grown to stationary phase in medium lacking uracil and containing 0.6% casamino acids, diluted 1:100 in the same medium and grown for 18 hr at 30°. Cell densities were normalized to OD_600_ before measuring fluorescence with a BioTek Synergy H1 microplate reader at an emission wavelength of 528 nm. The mean relative fluorescence units were calculated from three technical replicates for each biological replicate.

### Data Availability

Strains and plasmids are available upon request. The authors affirm that all data necessary for confirming the conclusions of the article are present within the article, figures, and supplemental material. Supplemental files are available at FigShare. Supplemental File S1 contains all supplemental tables and figures. Supplemental File S2 contains raw and validated SGA data. Supplemental File S3 contains R code used to analyze mass spectrometry and RNA sequencing count data. The gene expression data from the RNA sequencing analysis have been deposited in NCBI’s Gene Expression Omnibus (Edgar *et al.* 2002) and are accessible through GEO Series accession number GSE174145. The mass spectrometry proteomics data have been deposited to the ProteomeXchange Consortium via the PRIDE (Perez-Riverol *et al.* 2019) partner repository with the dataset identifier PXD025934.

## RESULTS

### Mistranslation by tRNA^Pro^_G3:U70_ and tRNA^Ser^_UCU, G26A_

Our goal was to compare the impact of two tRNA variants that mistranslate at a similar frequency, calculated from the ratio of mistranslated peptides to peptides containing the wild-type residue, but cause different amino acid substitutions. Previously, we have identified yeast tRNAs that mistranslate alanine or serine at different codons (Hoffman *et al.* 2017a; Berg *et al.* 2017, 2019b). Our quantification of their mistranslation frequencies using mass spectrometry led us to further analyze two tRNAs (Figure 1A). The first tRNA variant, tRNA^Pro^_G3:U70_, contains a G3:U70 base pair in the acceptor stem of a proline tRNA and mistranslates alanine at proline codons (Hoffman *et al.* 2017). The second tRNA variant, tRNA^Ser^_UCU, G26A_, is a serine tRNA with an arginine anticodon and a G26A secondary mutation to dampen tRNA function and mistranslates serine at arginine codons (Berg *et al.* 2017, 2019b). As one of the planned comparisons was a synthetic genetic array analysis, we constructed strains where the gene encoding the tRNA variants was integrated into the genome at the *HO* locus and selected with a clonNAT resistance marker. A third strain was created as a control with the clonNAT resistance marker alone integrated at the *HO* locus.

**Figure 1.**
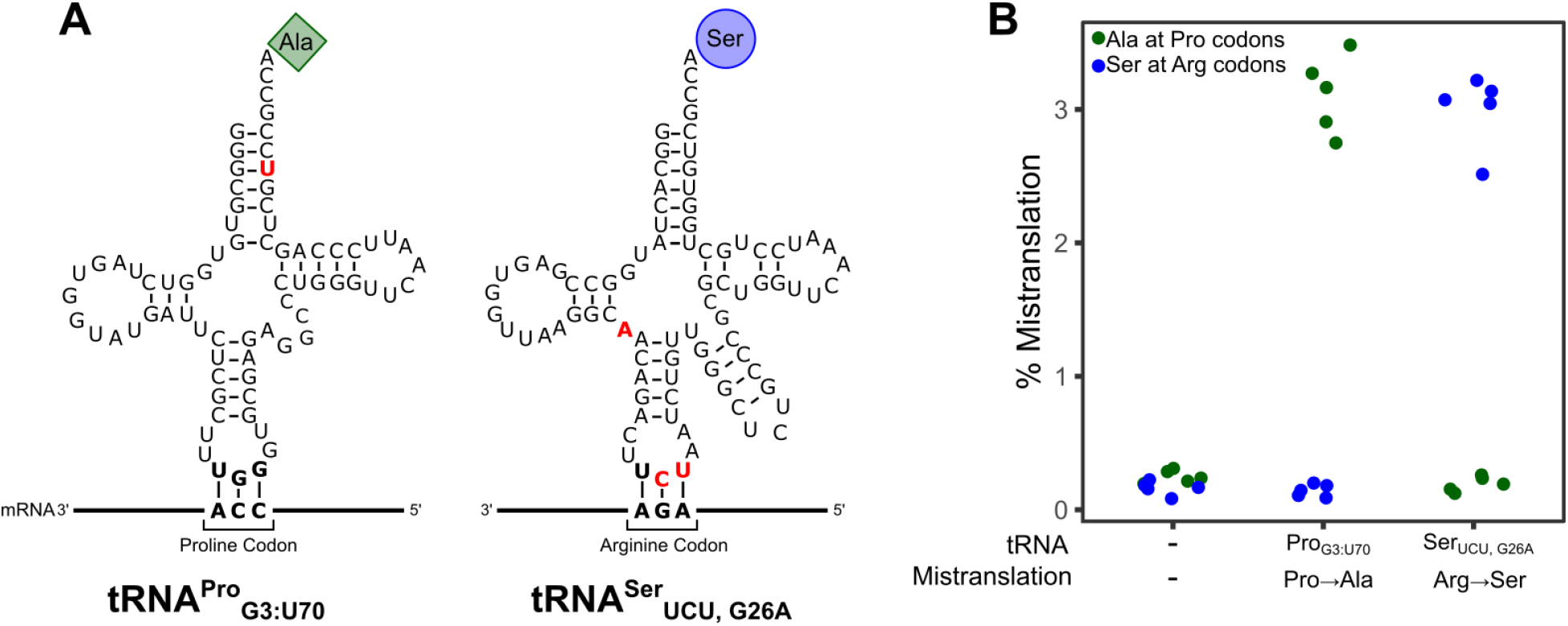
tRNA variants that mistranslate. **A.** Secondary structure of tRNA^Pro^_G3:U70_, which mistranslates alanine at proline codons, and tRNA^Ser^_UCU, G26A_, which mistranslates serine at arginine codons. Nucleotides colored in red denote differences compared to the wild-type tRNA^Pro^ or tRNA^Ser^, respectively. **B.** Mass spectrometry analysis of the cellular proteome was performed on a control strain with no additional tRNAs (CY8611) or strains expressing mistranslating tRNA variants tRNA^Pro^_G3:U70_ (CY8612) or tRNA^Ser^_UCU, G26A_ (CY8614). Each point represents one biological replicate. Each strain expressing a mistranslating tRNA had statistically higher frequency of mistranslation compared to the control strain (*p* ≤ 0.05; Welch’s *t*-test).

Using mass spectrometry, we determined the frequency of mistranslation for each strain (Figure 1B; Table S2). The frequency of proline to alanine mistranslation in the strain expressing tRNA^Pro^_G3:U70_ was 3.1%. Arginine to serine substitution in the strain expressing tRNA^Ser^_UCU, G26A_ was a similar 3.0%. The corresponding substitutions occurred at a frequency of approximately 0.2% in the control strain. Based on the mistranslated peptides identified from the proteomic mass spectrometry analysis of the proteome, tRNA^Pro^_G3:U70_ predominantly mistranslated alanine at CCA proline codons, whereas tRNA^Ser^_UCU, G26A_ decoded primarily AGA arginine codons (Figure S2).

### Characterizing the impact of mistranslation on cellular phenotypes

We determined the effect of the two tRNAs on cell growth in liquid culture under various nutrient conditions: yeast peptone (YP) or synthetic complete (SC) media with glucose or galactose as the carbon source, YP with glucose containing 4% ethanol, SC with monosodium glutamate and glucose as the nitrogen and carbon sources, respectively, and minimal media containing ammonium sulfate as the nitrogen source, 2% glucose, adenine, histidine, leucine, lysine and methionine. Both strains expressing mistranslating tRNAs grew slower than the control strain in all conditions (Figure 2A). These differences were statistically significant (*p* ≤ 0.05; Welch’s *t*-test) with the exception of the strain expressing tRNA^Pro^_G3:U70_ grown in medium containing 4% ethanol. In rich media (YPD), the alanine mistranslating tRNA (tRNA^Pro^_G3:U70_) increased doubling time to 80 min as compared to 74 min for the control strain. The serine mistranslating tRNA (tRNA^Ser^_UCU, G26A_) had a somewhat greater impact on growth, increasing doubling time to 85 min. This trend of the arginine to serine substitution having a greater effect was seen under all conditions, except in rich media containing galactose as the carbon source (YP + Galactose) where the two mistranslating tRNAs similarly increased doubling time relative to the control. Also indicative of the effect of environment on the phenotypic consequences of specific mistranslating tRNAs, the negative impact of tRNA^Ser^_UCU, G26A_ was greater when cells were grown in synthetic complete media with either glucose or galactose as the carbon source.

**Figure 2.**
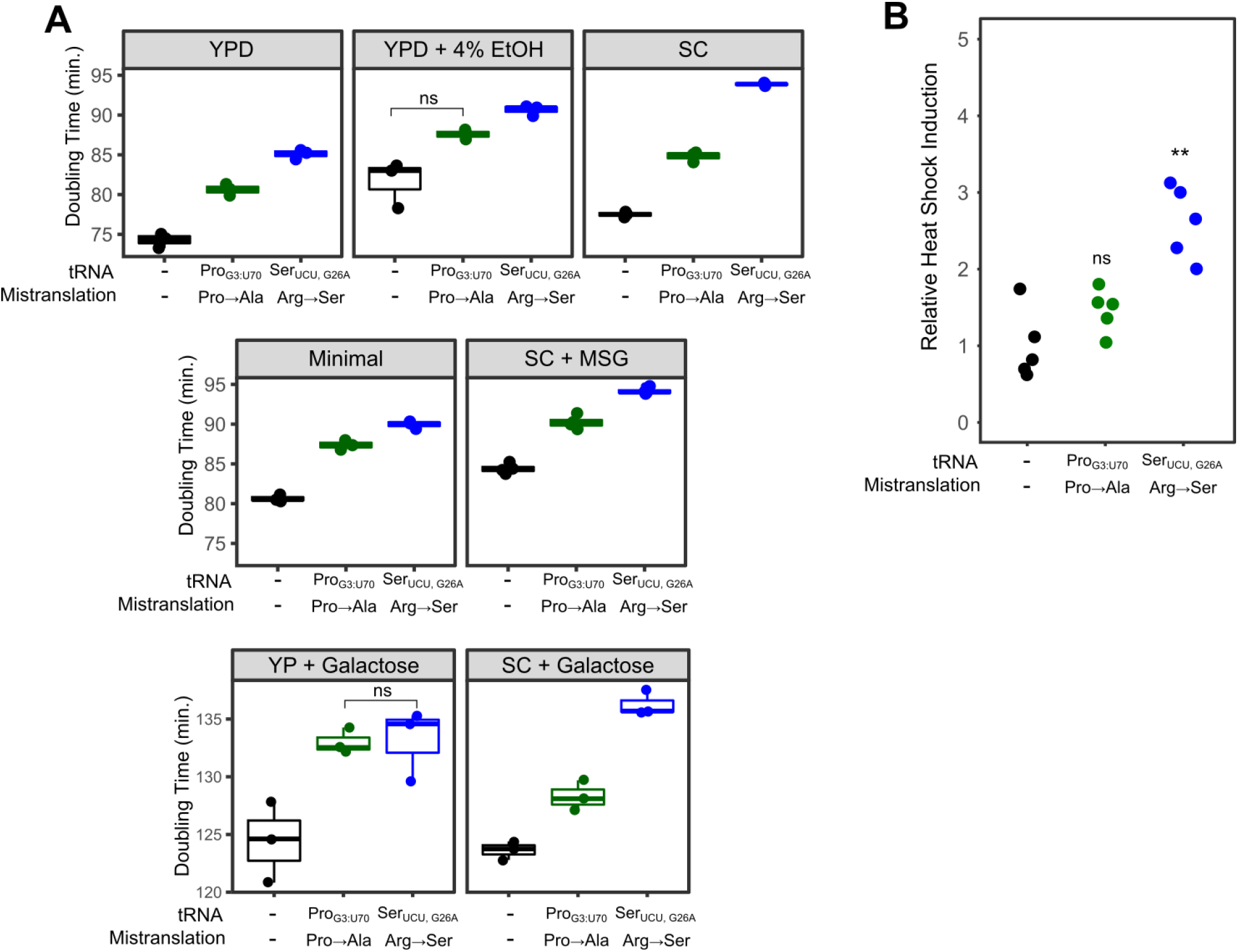
Phenotypic characterization of mistranslating strains. **A.** Growth rates for either a control strain with no additional tRNA (CY8611) or strains expressing mistranslating tRNA variants tRNA^Pro^_G3:U70_ (CY8612) or tRNA^Ser^_CU, G26A_ (CY8614) were determined from growth curves of the strains diluted to an OD_600_ ~ 0.1 in various media and grown for 24 hr. (YPD – yeast peptone with 2% glucose, YP + Galactose – yeast peptone with 2% galactose, SC – synthetic complete with ammonium sulfate as the nitrogen source and 2% glucose or 2% galactose as the carbon source, minimal – medium containing ammonium sulfate as the nitrogen source, 2% glucose and adenine, histidine, leucine, lysine and methionine, SC + MSG – synthetic complete media with monosodium glutamate and glucose as nitrogen and carbon sources). Doubling time was calculated with the R package “growthcurver” (Sprouffske and Wagner 2016). Each point represents one biological replicate. All comparisons within a growth condition are statistically different (*p* ≤ 0.05; Welch’s *t*-test), except where indicated (ns =not significant). **B.** Strains described in A were transformed with a GFP reporter transcribed from a promoter containing heat shock response elements, grown to saturation in media lacking uracil, diluted 1:100 in the same media and grown for 18 hours at 30°. Cell densities were normalized and fluorescence measured. Each point represents one biological replicate. Statistical comparisons were made between strains expressing a variant tRNA and the control strain (ns =not significant, ** *p* ≤ 0.005; Welch’s *t*-test).

Mistranslation results in proteotoxic stress and a heat shock response (Berg *et al.* 2017). To determine the level of heat shock response found in the strains containing the tRNA variants, we measured fluorescence arising from GFP under the control of a synthetic heat shock promoter (HSE; Brandman *et al.* 2012; Figure 2B). Expression of HSE-GFP was 2.8-fold greater in cells containing tRNA^Ser^_UCU, G26A_ (Arg→Ser) than in the control strain. In contrast, expression of HSE-GFP in tRNA^Pro^_G3:U70_ (Pro→Ala) was not statistically different from the control.

### Mistranslating tRNAs have different genetic interactions

We used a synthetic genetic array (SGA) analysis to identify the genetic interactions of the two tRNAs and provide another comparison of the impact of the substitutions. Preliminary screens of the strain expressing tRNA^Pro^_G3:U70_ identified a greater percentage of interactions with the temperature sensitive collection than the deletion collection, therefore the former was chosen for the comparison of tRNA^Pro^_G3:U70_ and tRNA^Ser^_UCU, G26A_.

SGA analyses were performed in parallel for the mistranslating and control strains. Of the 1016 alleles in the temperature sensitive collection, 18 had a negative genetic interaction with tRNA^Pro^_G3:U70_ (Pro→Ala), whereas 78 alleles had a negative interaction with tRNA^Ser^_UCU, G26A_ (Arg→Ser). No positive genetic interactions were identified. Genetic interactions were validated by remaking the double mutant strains, spotting normalized densities of the double mutants and their control strain on selective medium and measuring growth after two days. After validation, 10 and 47 alleles showed negative genetic interactions with tRNA^Pro^_G3:U70_ and tRNA^Ser^_UCU, G26A_ respectively (Figure 3). The increased number of genes identified with tRNA^Ser^_UCU, G26A_ parallels its greater effect on growth and greater heat shock response. Raw data and validated genes are found in Supplemental File S2.

**Figure 3.**
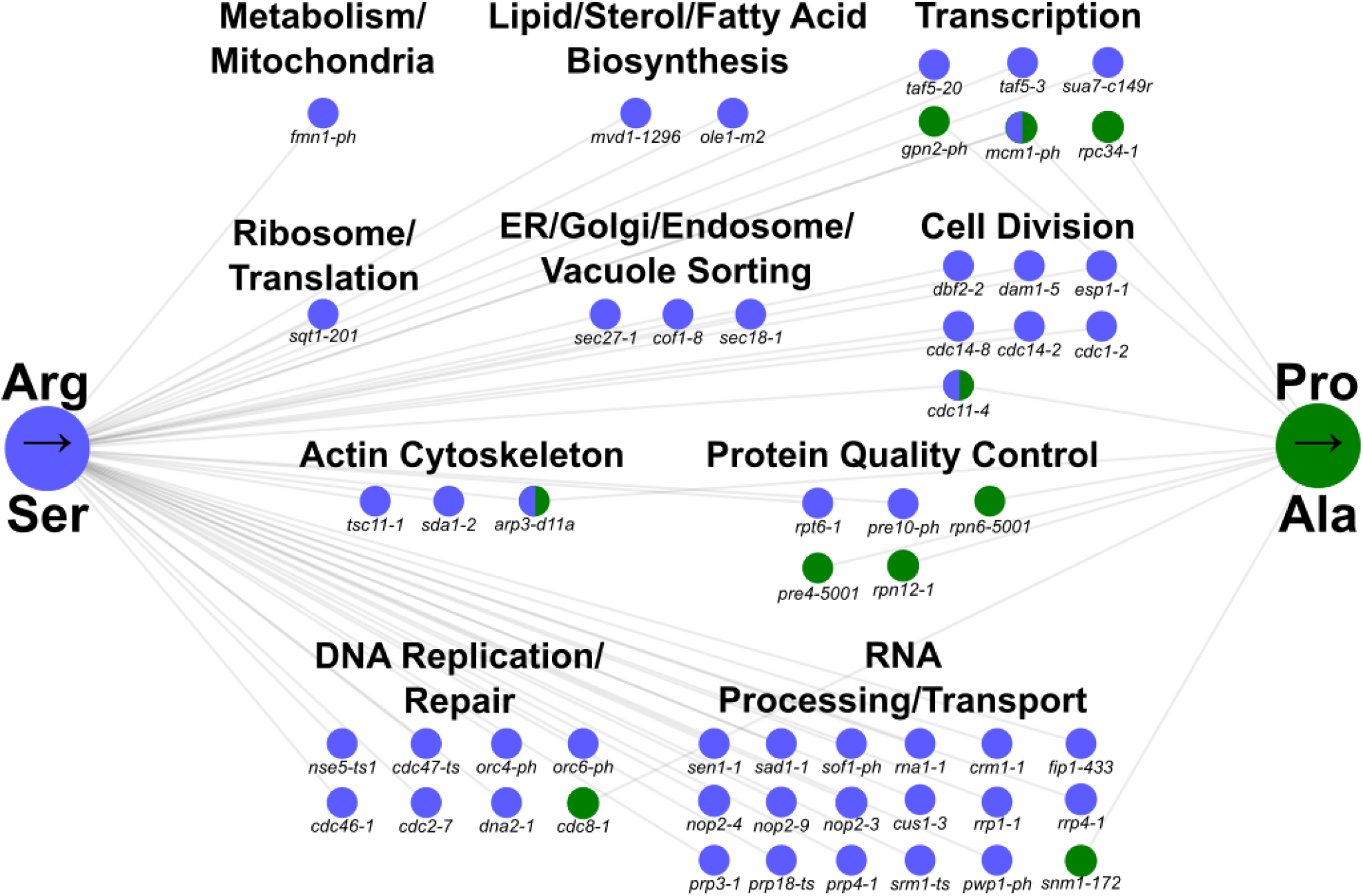
Negative genetic interaction networks of the mistranslating tRNAs. Genetic interaction network of temperature sensitive alleles that have negative genetic interactions with tRNA^Pro^_G3:U70_ (green) and tRNA^Ser^_UCU, G26A_ (blue). Nodes represent alleles and edges represent negative genetic interactions.

Analysis of the biological function of the genes displaying negative genetic interactions with the mistranslating tRNA variants revealed both distinct and common processes. Not surprisingly, many of the processes were also identified in chemical-genetic screens with amino acid analogs that are incorporated into protein (Berg et al. 2020). As mentioned above, the most notable difference in the genetic interaction patterns between the two tRNAs is the increased number with tRNA^Ser^_UCU, G26A_. The difference in number of interactions makes evaluating whether there is enrichment for one of the tRNAs difficult. In general, both tRNAs have genetic interactions with genes involved in transcription and protein quality control. The former suggests a potential importance for a transcriptional response to mistranslation, while the later highlights the role of proteasomal components in coping with mis-made proteins (Shcherbakov et al. 2019). While a negative genetic interaction with a mistranslating tRNA variant may indicate the gene/process is required for cells to cope with the resulting mis-made protein, it may also indicate that the process is sensitive to specific mistranslation. For example, 15 of the 16 genes involved in RNA processing and transport were specifically found with tRNA^Ser^_UCU, G26A_ suggesting the possibility that the individual proteins or the process may be particularly sensitive to substitution at arginine codons. The RNA processing/transport genes provide an example that highlights that although processes were shared, specific genetic interactions differed; only three genes were synthetic with both tRNAs. This led us to evaluate whether amino acid composition of the encoded protein contributed to a gene being synthetic with a mistranslating tRNA, using the RNA processing/transport genes as a test case. The average arginine content of the proteins whose genes are synthetic with tRNA^Ser^_UCU, G26A_ varies from 7.8% (Sof1) to 2.5% (Rna1) as compared to the cellular average of 4.4% (Table S3). In several cases, the arginine content of the tRNA^Ser^_UCU, G26A_ interactors is less than the proline content (Sad1, Rna1, Fip1, Pwp1 and Srm1). This indicates that amino acid composition of the encoded proteins alone does not explain the specific negative genetic interactions with these genes.

### Impact of tRNA^Pro^_G3:U70_ or tRNA^Ser^_UCU, G26A_ on the transcriptome

We next compared the transcriptome profiles of strains expressing tRNA^Pro^_G3:U70_ or tRNA^Ser^_UCU, G26A_. To ensure that the strains had the identical genetic background without possible suppressor mutations, we transformed plasmids expressing the mistranslating tRNAs or empty plasmid into yeast strain BY4742. The transcriptome of each strain was profiled using an RNA sequencing approach. Principal component analysis (PCA) of the three transcriptome profiles demonstrated that ~ 45% of the variance could be explained in two components (Figure 4A). The wild-type control strain clusters separately from the strains expressing mistranslating tRNAs in the first component and the two strains expressing different mistranslating tRNAs separate in the second component. This suggests that mistranslation of alanine at proline codons initiates a distinct response from mistranslation of serine at arginine codons.

**Figure 4.**
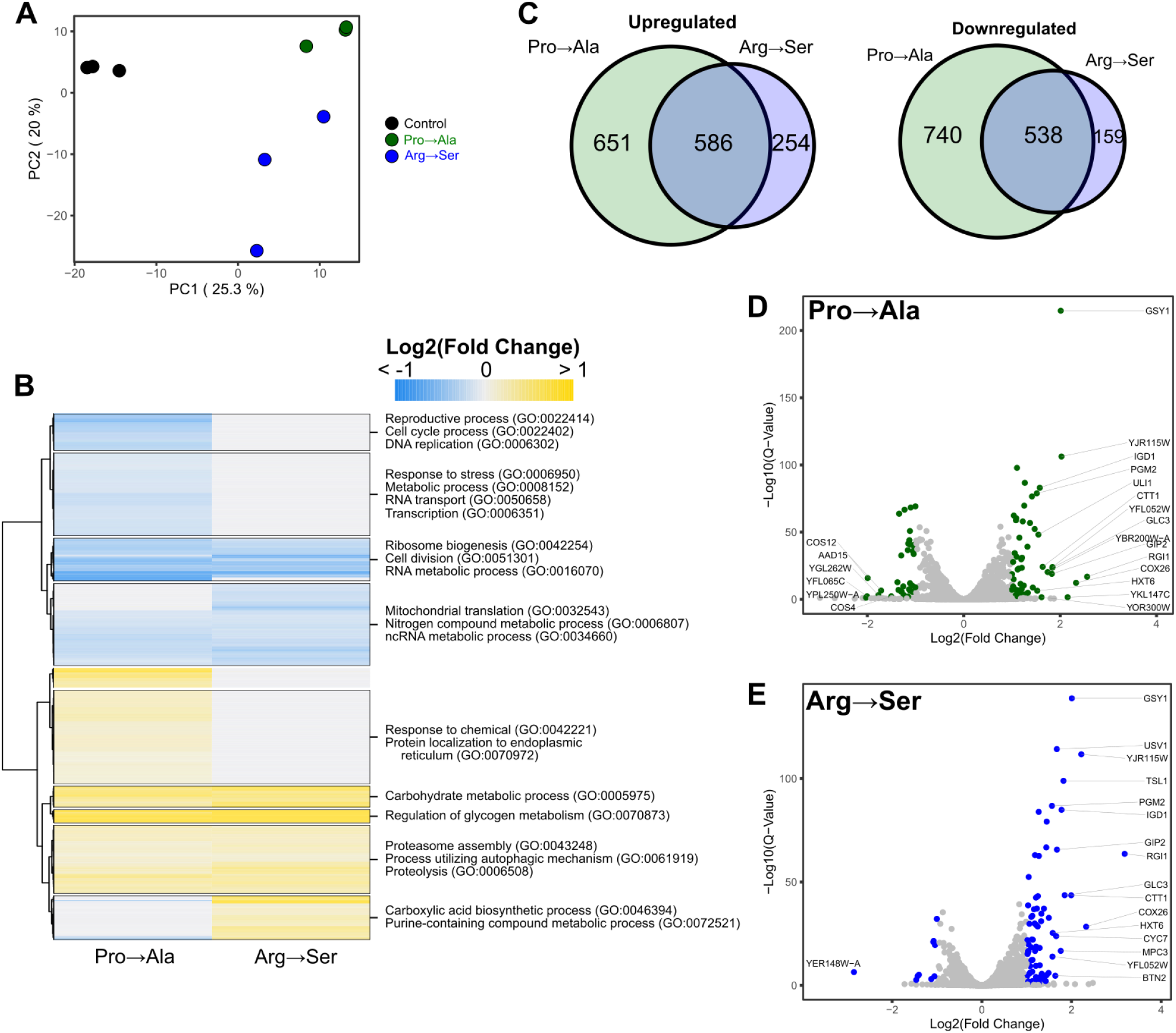
Transcriptome analysis of strains expressing mistranslating tRNA variants. **A.** Principal component analysis of centered log ratio normalized reads from BY4742 expressing either tRNAProG3:U70 (Pro→Ala), tRNASerUCU, G26A (Arg→Ser) or an empty vector (WT). Each point represents one biological replicate (n =3). **B.** Heatmap of hierarchical clustered differentially expressed genes (*p* ≤ 0.05) in response to mistranslation. Fold-change for each gene is the average from three biological replicates. Upregulated genes are colored yellow while downregulated genes are colored blue. Significantly enriched GO terms within each cluster are annotated. **C.** Venn diagram of upregulated and downregulated genes (*p* ≤ 0.05) in yeast strain BY4742 transformed with a centromeric plasmid expressing tRNAProG3:U70 (Pro→Ala) or tRNASerUCU, G26A (Arg→Ser) as compared to BY4742 transformed with empty vector. **D.** Volcano plot highlighting in green differentially expressed genes with > 2-fold changes in tRNAProG3:U70 relative to the control strain. Points with gene name labels have the largest fold-change relative to control. **E.** Volcano plot highlighting in blue differentially expressed genes with > 2-fold changes in tRNASerUCU, G26A as in D.

We then analyzed the differentially expressed genes in each mistranslating strain as compared to the control strain. A total of 2897 genes (44.7%) were differentially expressed in at least one strain (Benjamini-Hochberg adjusted *p*-value cut off ≤ 0.05). A heatmap of the fold-change in transcript levels for these genes is shown in Figure 4B. A Venn diagram of the overlap of up- and downregulated genes caused by the mistranslating tRNAs is shown in Figure 4C. Of note, distinct groups of genes with unique responses based on the type of mistranslation were identified. For example, genes involved in cell cycle, DNA replication, transcription and response to stress were down-regulated predominantly in the strain expressing tRNA^Pro^_G3:U70_ (Pro→Ala) whereas genes involved in response to chemical and protein localization to the endoplasmic reticulum were upregulated. Genes involved in carboxylic acid biosynthetic processes and purine metabolism were upregulated predominantly in the strain expressing tRNA^Ser^_UCU, G26A_ (Arg→Ser). There were also common sets of genes differentially expressed in both mistranslating strains. For example, genes involved in mitochondrial translation, cell division, ribosome biogenesis and RNA metabolism were downregulated, and genes involved in carbohydrate metabolism and proteolysis were upregulated in both strains. Overall, these results demonstrate that while a common set of genes respond to mistranslation, each of the two mistranslating tRNAs induce a unique transcriptional response.

A more stringent analysis of the transcriptional changes, considering only genes with a ≥ 2-fold change in expression, revealed 55 and 78 genes upregulated and 34 and 71 genes down-regulated in strains expressing tRNA^Pro^_G3:U70_ and tRNA^Ser^_UCU, G26A_, respectively (Figure 4D, E). Amongst the upregulated genes, 36 are common to both mistranslating tRNA variants. These included genes involved in carbohydrate metabolism, mainly glycogen and trehalose biosynthesis (e.g. *GLC3, IGD1, GSY1, GSY2, MRK1, PGM2, PIG2, GIP2, TPS2, TPS2, TSL1*). The upregulation of genes involved in the synthesis of these storage carbohydrates is consistent with previous studies demonstrating that glycogen and trehalose accumulate under the stress of nutrient starvation or heat shock (Lillie and Pringle 1980; Hottiger *et al.* 1987; Parrou *et al.* 1997) and that trehalose stabilizes proteins and suppresses aggregation (Singer and Lindquist 1998). Genes involved in protein folding were also upregulated in both strains. These include the small heat shock protein, *HSP42*, which suppresses misfolded protein aggregation, components of the chaperonin complex (*TCP1)* and genes that direct ubiquitination of misfolded proteins (*ROQ1)*. Five genes (*FET3, AAD15, GFD2, YOR338W and ALD6*) were downregulated in both strains expressing mistranslating tRNA variants.

## DISCUSSION

Mistranslation can generate many distinct substitutions that differ with regard to the amino acid (or stop codon) replaced and the amino acid that is inserted. Previously, we found that the frequency of mistranslation resulting from tRNA variant impacts the physiological consequences with increasing frequencies of mistranslation having more severe effects (Berg et al. 2019b). Here we show that two tRNA variants that mistranslate different codons with a frequency of ~3% elicit different effects on growth, heat shock response and the transcriptome and have distinct sets of genetic interactions.

### Significance of the differing impacts of tRNA variants

tRNA variants with the potential to mistranslate are found in the human population (Berg *et al.* 2019a). Approximately 20% of all individuals have either an anticodon mutation in tRNA^Ser^ or tRNA^Ala^ or an acceptor stem mutation that generates a G3:U70 base pair, the major identity element for charging with alanine (Hou and Schimmel 1988; Francklyn and Schimmel 1989). The ability of mutations in other parts of the tRNA, such as the Darm (Hirsh 1971), to alter decoding specificity suggests that more mistranslating variants may exist amongst the more than 600 human tRNA encoding genes. We hypothesize that mistranslating tRNAs are genetic modifiers of disease by increasing the level of proteotoxic stress. This increased stress could lead to earlier onset and/or increased severity of diseases that have proteotoxic stress as their hallmark, including many neurodegenerative diseases, cardiomyopathies and hearing loss (Willis and Patterson 2013; Labbadia and Morimoto 2015; McLendon and Robbins 2015; Jongkamonwiwat *et al.* 2020). In this framework, our results suggest that even for tRNA variants with a similar mistranslation frequency, their impact on disease would differ based on the specific substitution. For example, because of the increased heat shock observed for tRNA^Ser^_UCU, G26A_ (Arg→Ser), we expect tRNA^Ser^_UCU, G26A_ to have a greater impact than tRNA^Pro^_G3:U70_ (Pro→Ala). Moreover, given that diseases often result from one genetic change or a polygenic mix of genetic changes, the finding that mistranslating tRNAs show unique patterns of negative synergy with specific genes is a strong indication that tRNA variants could act as genetic modifiers of distinct diseases. Synergy between a tRNA variant and a cellular mutation can act in either direction. Most commonly the mistranslating tRNA will alter the severity of a mutation in another gene. However, if a gene regulates tRNA expression or function, its mutation could change the impact of the mistranslating tRNA variant.

### Factors affecting the impact of mistranslating tRNAs

Before addressing what alters the impact of a mistranslating tRNA, it is important to consider how proteome-wide mistranslation could affect a cell. First, as indicated above, mistranslation results in protein mis-folding and aggregation that in turn give rise to proteotoxic stress. On its own this will impact cell function. Second, decreased expression of individual cellular proteins often has a phenotypic consequence (Breslow *et al.* 2008). Mistranslation acts at the proteome level to decrease the amount of each protein that is functional, effectively creating hypomorphs. Global mistranslation may exaggerate the effect because of the potential that multiple genes in a pathway as well as those in redundant pathways can be compromised.

Given equal frequencies of substitution, the relative impact of two mistranslating tRNAs is determined by the absolute number of amino acid changes, the chemical nature of the amino acid change and the structural context of the target amino acid in different proteins. Variants that mistranslate amino acids found more abundantly in proteins have more targets for mis-incorporations and thus a greater probability of inducing protein unfolding and/or affecting a hypomorphic gene. Added to this, one also needs to take into account that a specific mistranslating tRNA has the potential to affect some proteins more than others because amino acid composition varies from protein to protein. Furthermore, the extent an amino acid is used depends on two aspects: the abundance of codons for the amino acid in the genome and the level of expression of those proteins containing the amino acid. The combination of these factors makes predicting the absolute number of amino acid substitutions difficult, especially since protein expression and regulation are dynamic. Related to this, it was interesting to observe the extent to which the growth media affected the relative impact of the mistranslating tRNAs. This can be explained by the fact that environmental conditions alter the expression of the genome and resulting proteome (Navarrete-Perea *et al.* 2021).

The properties of the amino acids being substituted is the second major factor affecting the impact of proteome wide mistranslation. In theory, more chemically diverse substitutions should have more profound effects on protein structure and function. In fact, the genetic code is predicted to have evolved by minimizing the impact of mutations on the resulting change in these properties (Koonin and Novozhilov 2017). Woese *et al.* (1966) proposed the polar requirement (PR) scale for the relatedness of amino acids. The PR scale is based on the solubility of each amino acid in pyridine: proline, alanine, serine and arginine score 6.6, 7.0, 7.5 and 9.1, respectively. For mistranslating tRNAs, the substitution would be expected to be less severe if the amino acids are close on the PR scale. Fitting with this we see less impact for Pro→Ala than Arg→Ser. The distinct biochemical properties of each amino acid may also confer special functions to a protein that are lost or gained upon substitution; for example, the phosphorylation of serine and the prevalence of arginine in nuclear localization signals. We considered substitution matrices as another way to evaluate the impact of mistranslation. Substitution matrices describe the likelihood that one amino acid would be substituted for another over evolutionary time. Accepted point mutation (PAM) matrices extrapolate from alignments of closely related sequences (Dayhoff and Eck 1968), whereas blocks substitution matrix (BLOSUM) reflects changes found in more distantly related proteins (Henikoff and Henikoff 2000). We found that the substitution scores in these matrices do not match the magnitude of physiological effect seen for the substitutions tested here. This may be because the matrices are compiled from evolutionarily related protein sequences where certain substitutions are less likely to occur because of genetic code constraints, rather than functional properties of the protein.

We note that we have restricted our discussion to the roles of tRNAs in translation. In mammalian cells, tRNAs regulate many processes, either as intact molecules or fragments (Berg and Brandl 2020). It is possible that mistranslating tRNA variants affect biological processes in a translation independent manner.

### Mistranslating alanine at proline codons with tRNA^Pro^_G3:U70_

We first identified alleles encoding tRNA^Pro^_G3:U70_ (Pro→Ala) as spontaneous suppressors of a conditional *tti2-L187P* allele (Hoffman *et al.* 2017). The suppressor tRNA inserts alanine at the proline codon at position 187 in *tti2*, which restores nearly full function to Tti2. Mistranslation is due to a single base change of C70 to T in a tRNA^Pro^_UGG_ encoding gene. In the *tti2*-based selection, we identified four independent tRNA^Pro^_UGG_ genes with this mutation and estimate that it occurs at a frequency of approximately 10^-6^ in yeast populations. tRNAs with G3:C70 base pairs, a single mutation removed from creating the major G3:U70 alanine identity element, are found in bacteria, archaea and yeast species, but are rare in eukaryotes other than yeast (Berg *et al.* 2018). tRNA^Lys^_G3:U70_ variants have been isolated from *E. coli* (see Prather *et al.* 1984). Combined with the prevalence of tRNA^Pro^_G3:U70_ in yeast populations, these findings suggest that in single cell organisms the ability to mistranslate codons as alanine may provide an advantage to the population. The absence of G3:C70 in most eukaryotic tRNAs suggests that the mistranslation resulting from a change to U70 would be deleterious for these species.

In summary, mistranslating tRNAs impact cells in ways that are specific to the type of mistranslation. The consequences of these mistranslating tRNAs will in turn depend upon the genotype of the organism and the environment. It will be important to consider these differences when analyzing tRNAs as genetic modifiers of disease.

## ACKNOWLEDGMENTS

We would like to thank Ricard Rodriguez-Mias for assisting with the mass spectrometry and maintaining the instruments, Mario Leutert for assistance with the R2P1 sample preparation and data collection, Stephanie Zimmerman for providing the code to analyze codon distribution in mistranslated peptides, Onn Brandman for the HSE-GFP reporter plasmid, Brenda Andrews for yeast strain Y7092 and Ecaterina Cozma for critical reading of the manuscript.

